# Discovery of specific activity of 2-HPA acting on the membrane progestin receptor α (paqr7) by purification of natural products from the marine algae *Padina*

**DOI:** 10.1101/2023.09.12.557470

**Authors:** Mohammad Tohidul Amin, Mrityunjoy Acharjee, Md. Maisum Sarwar Jyoti, Md. Rezanujjaman, Md. Maksudul Hassan, Md. Forhad Hossain, Saokat Ahamed, Shinya Kodani, Toshinobu Tokumoto

## Abstract

Membrane progestin receptors (mPRs) are members of the progestin and adipoQ (PAQR) receptor family that are stimulated by endogenous steroids to initiate rapid intracellular signalling through a nongenomic pathway. Previously, water-soluble compounds with mPRα-binding activity from the marine algae *Padina arborescens* were fractionated by HPLC steps. In this study, the structure of one of the major compounds in the fraction was identified as 2-hydroxypentanoic acid (2-HPA) using Nuclear Magnetic Resonance spectroscopy. 2-HPA showed a substantial competitive binding affinity for hmPRα in the GQD-hmPRα binding assay. In contrast, synthetic structural analogues of 2-HPA showed no competitive binding activity. The physiological activity of 2-HPA and its analogues was then investigated using *in vitro* goldfish and *in vivo* zebrafish oocyte maturation and ovulation assays. As with the hmPRα binding assay, only 2-HPA showed inhibitory activity on oocyte maturation and ovulation of fish oocytes. Furthermore, the inhibitory activity of 2-HPA was compared between S- and R-type 2-HPA. The results showed that both types had the same level of activity. These results indicate that 2-HPA, found as a secreted compound from *Padina arborescens*, is a novel mPRα antagonist and its chemical structure is highly restricted to show its activity.

## Introduction

Membrane progestin receptors (mPRs) are seven-or eight-transmembrane plasma membrane receptors for progesterone or its analogues. mPRs are members of a novel family of progestin and adipoQ receptors (PAQRs) consisting of 11 genes that show homology to the adipoQ receptors^1^. The mPR molecule has five types, α, β, γ, δ and ε, corresponding to PAQR7, PAQR8, PAQR5, PAQR6 and PAQR9, respectively^2,3^. It is also known that mPRs are conserved in a wide range of vertebrate cells, from humans to fish, and are expressed in a variety of tissues, including brain and kidney^2,4^. To date, mPRs have been implicated in biological regulation, including oocyte maturation in fish and amphibians, induction of reproductive behaviour in mammals^5-8^. Recently, it has attracted particular attention for its involvement in brain development as a target for neurosteroids^9^. In addition, mPRs have long been shown to be highly expressed in cancer cells and are thought to mediate the pathways of cancer growth and invasiveness^10,11^. This group of mPR molecules is known to be involved in the regulation of many cellular activities, and the search for mPR reactive substances that respond to them is ongoing^12,13^. In the initial studies of mPR discovery, Org OD 02 was already discovered as an mPR-specific agonist and has been used for research purposes^11,14^. We have also shown that Org has the ability to induce oocyte maturation in fish^15^. The experimental systems for *in vitro* oocyte maturation in fish and amphibians have long been shown to be induced by nongenomic actions of progesterone, and were have been used as a test for nongenomic actions, leading to the discovery of mPR^5^. This assay has been used exclusively as a method for testing mPR reactive substances^16,17^.

We have developed *in vivo* zebrafish oocyte maturation and ovulation induction methods as screening methods for mPR-responsive substances, as well as a test method using the modified luciferase gene (Glosensor) to measure intracellular cAMP concentration^18,19^. We have also succeeded in expressing and purifying mPRα molecules using yeast, and have developed a high-throughput screening method for mPR-responsive substances using GQD-mPRα, which is a nanoparticle-associated mPRα^20,21^. On the other hand, we are also trying to isolate and purify mPR-reactive natural products using these screening methods^22^. We have discovered the presence of mPR-reactive substances in the seawater of coral reefs in Mauritius and recently succeeded in separating them into two peaks^23,24^. In this study, the chemical structure of the main component of one of these two peaks was successfully determined. The results of the physiological activity of the resulting organic component showed that it is an inhibitor of oocyte maturation, i.e., a novel antagonist of mPRα. mPRα-specific antagonists have never been reported before and are expected to be useful research tools and novel drug candidates in mPR research in the future. It is expected to be a useful research tool in mPR research and a new drug candidate in the future.

## Results

In a previous study, compounds with mPRα-binding activity from secretion material of the marine algae *Padina arborescens* were fractionated as two peaks by HPLC steps. Compounds in both of two peaks (Peak1 and Peak2) showed competitive binding activity against human mPRα and inhibitory activity on fish oocyte maturation and ovulation. In this study, we tried to identify the chemical structure of the active compound by chemical analysis. The Peak2 was subjected to NMR analyses using ^1^H-NMR, ^13^C-NMR, DQF-COSY, TOCSY, HMQC and HMBC (Figures S1-S6). Analysis of the NMR data indicated that Peak2 was a mixture of several compounds. As a result of the analysis of the 2D NMR data, the main component was determined to have the carbon skeleton of 2-hydroxypentanoic acid (2-HPA) with assignment of chemical shift values (Figure 1) (Table. S1). Briefly, TOCSY and DQF-COSY indicated the proton spin system from position 2 to 5 (bold line in Figure 1). The characteristic chemical shifts (δH 4.06, δC 64.7) of position 2 indicated the presence of a hydroxy residue. The HMBC correlation from the proton at position 2 to the carbon at position 1 suggested a carboxy residue at position 1. Although the material (Peak2) was a mixture of several compounds, we proposed that the carbon skeleton structure of 2-HPA may be essential for the activity.

**Figure 1.**
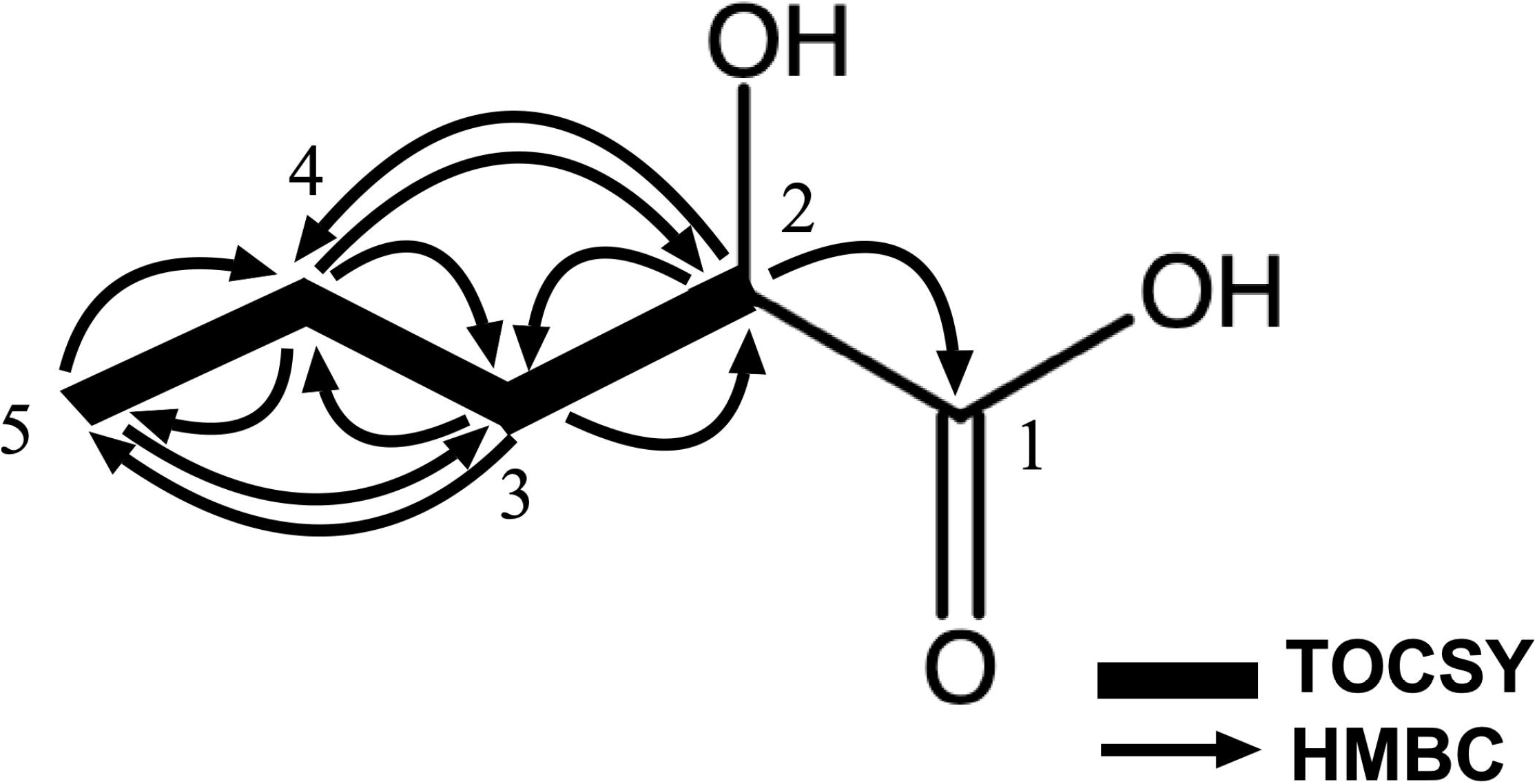
Key correlations of 2D NMR spectra of 2-hydroxypentanoic acid. Numbers indicate carbon position. One-sided arrow indicates HMBC correlation. Bold line indicates TOCSY correlation.

Subsequently, the mPRα-binding activity of 2-HPA and its analogues was evaluated using the newly established GQD-hmPRα binding assay, which allows high through put screening of mPR-interacting compounds. Closely related 2-HPA analogues were selected as follows: hydroxy group missing version of 2-HPA; valeric acid, hydroxy group position altered version; 3-hydroxyvaleric acid, shorter fatty acid chain; 2-hydroxybutyric acid, longer fatty acid chain; 2-hydroxyhexanoic acid, missing double bonded oxygen; 1,2pentanediol (Figure 2). As shown in Figure 3, the peak2 fraction isolated from *Padina* (Peak2) and pure (S)-2-HPA reduced the fluorescence intensity in a concentration-dependent manner as comparable to the positive control, progesterone. The result indicated that 2-HPA possessed hmPRα-interacting activity as the Peak 2 fraction. In contrast, no decrease in fluorescence intensity was observed for analogues of 2-HPA such as the negative control estradiol. Among the 2-HPA and its analogues tested in this study, only 2-HPA has hmPR binding properties, suggesting that the binding of 2-HPA to the hmPR is specific and that the interaction is conformationally restricted.

**Figure 2.**
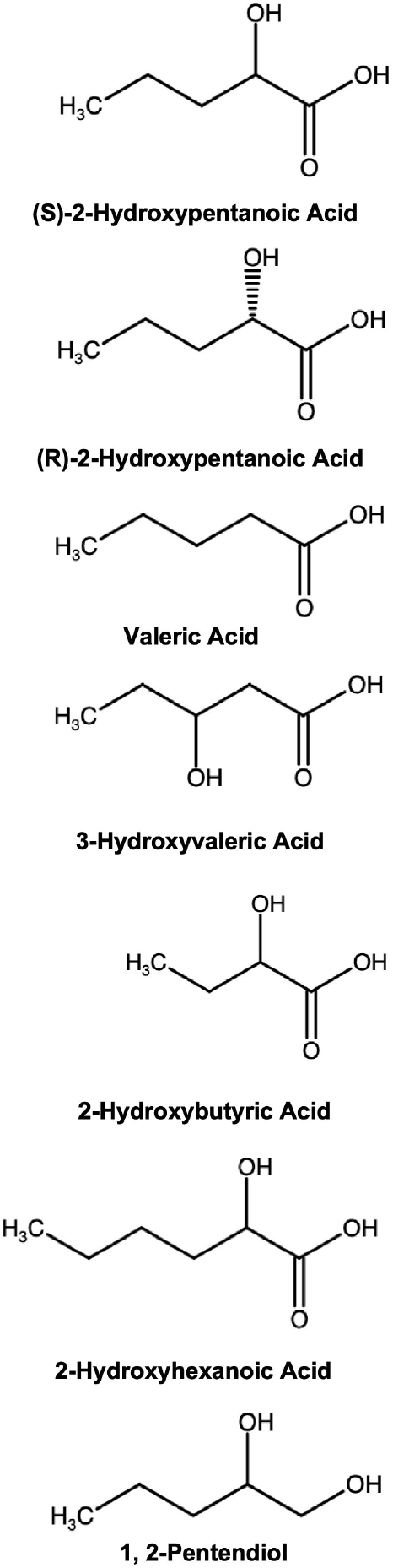
Structure of 2-HPA isomers and its analogues. Chemical structures of two isomers of 2-HPA and its analogues used in this study. Chemical structures were drawn by the Swiss target Prediction (http://www.swisstargetprediction.ch/).

**Figure 3.**
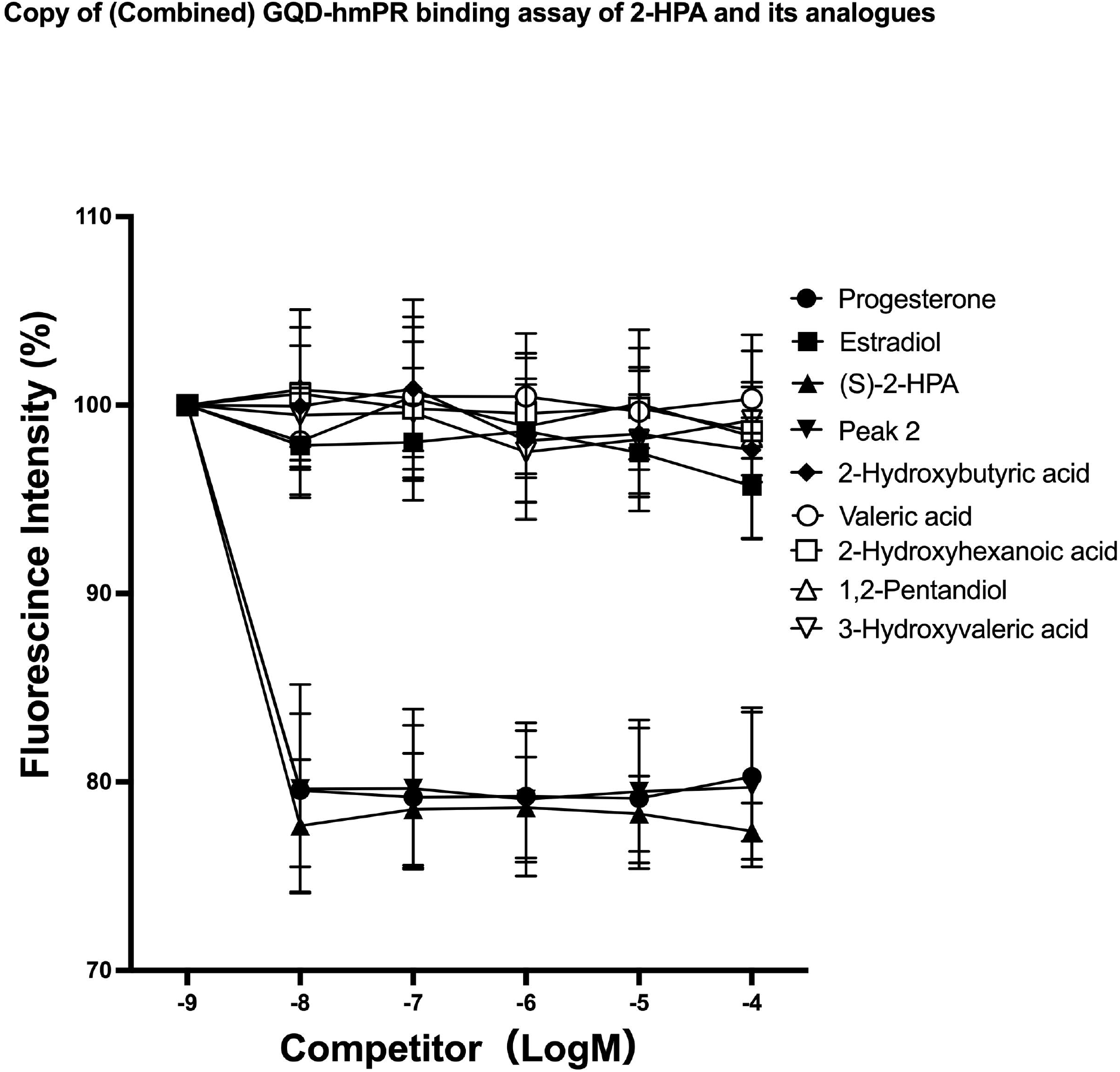
GQD-hmPRα binding assay of 2-HPA and its analogues. (A) Competition of the binding of P4-BSA-FITC with GQD-hmPRα by 2-HPA and its analogues. The dose-dependent effects of steroids (progesterone, estradiol-17β), (S)-type of 2-HPA, purified fraction from Padina (Padina-C) and 2-HPA analogues (see Figure 2) were studied.

The physiological activities of 2-HPA and its analogues were evaluated by an *in vitro* oocyte maturation assay using fish oocytes. Induction of oocyte meiotic maturation has been shown to be induced by mPRα mediated signal transduction through nongenomic action as a biological process made a discovery of mPRα^4^. Goldfish oocytes were used for large-scale analysis. 17α,20β-dihydroxy-4-prognen-3-one (DHP), is a natural agonist for the mPRs and induces oocyte maturation by binding to the mPRs on the cell surface of the oocyte.

Although no agonistic activity to induce oocyte maturation was observed when incubated compounds alone with oocytes (data not shown), 2-HPA showed antagonistic activity against DHP-induced oocyte maturation (Figure 4). As expected, only 2-HPA showed inhibitory activity on DHP-induced oocyte maturation and its analogues did not show any activity (Figure 4). Physiological activity of compounds was further confirmed by *in vivo* oocyte maturation and ovulation assay using zebrafish. Again only 2-HPA showed inhibitory activity on oocyte maturation and ovulation (Figure 6). Finally, the activity of 2-HPA isomers were compared. The results in Figure 6 show that both (R)-2-HPA and (S)-2-HPA inhibit oocyte maturation and ovulation at same magnitude of activity (Figure 5), with (R)-2 -HPA and (S)-2-HPA, strongly suggesting that both isomers act as antagonists for mPRα.

**Figure 4.**
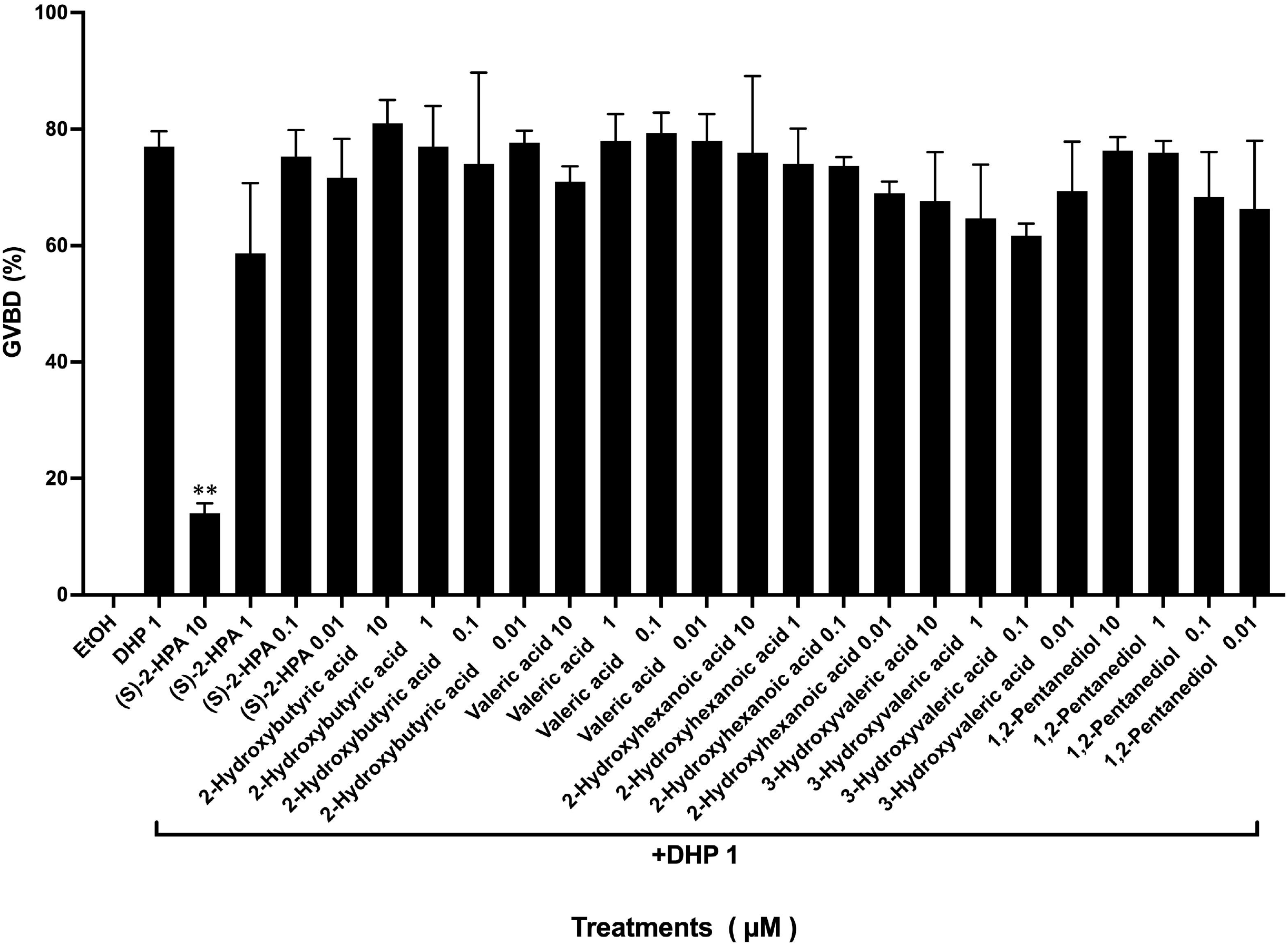
*In vitro* oocyte maturation assay of 2-HPA and its analogues. 2-HPA and its analogues were added to indicate concentrations, and then maturation was induced by 1 μM of 17,20β-DHP (+DHP 1). After six hours incubation, oocytes with or without germinal vesicle breakdown (GVBD) were counted and the percentage of GVBD was calculated. As a negative control, oocytes were incubated with 0.1 % ethanol (EtOH) or 1 μM of DHP alone as a positive control (DHP 1). The assay was performed in triplicate and the averaged percentage of GVBD is expressed as standard deviation.

**Figure 5.**
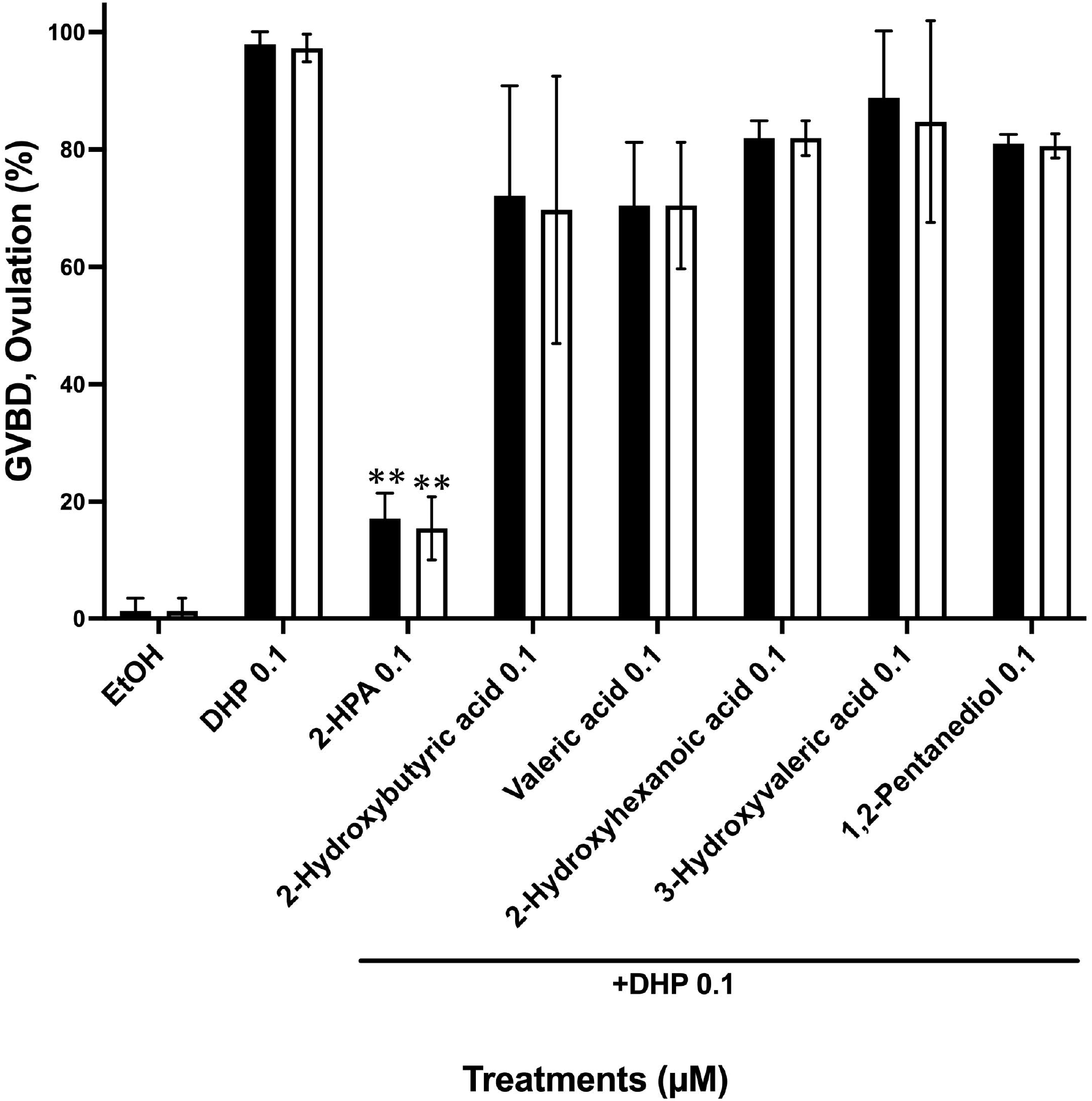
*In vivo* oocyte maturation and ovulation assay of 2-HPA and its analogues. The antagonistic activity of 2-HPA and its analogues against oocyte maturation and ovulation induction was analyzed by *in vivo* treatment in zebrafish. 2-HPA and its analogues were added into the water at 0.1 μM, and then maturation and ovulation were induced by the addition of 0.1 μM of 17,20β-DHP (DHP 0.1). After four hours of treatment with compounds by addition to water, %GVBD (closed column) and %ovulation (open column) were determined by scoring the oocytes that had become transparent and formed an egg membrane by egg activation. Fish were incubated with 0.01% ethanol (EtOH) as a negative control or with 0.1 μM DHP alone as a positive control (DHP 0.1). Three fish were used per treatment. Means of data are presented with standard deviation. Asterisks represent significant differences between DHP alone and DHP with 2-HPA treatment (** P≤m 0.001).

**Figure 6.**
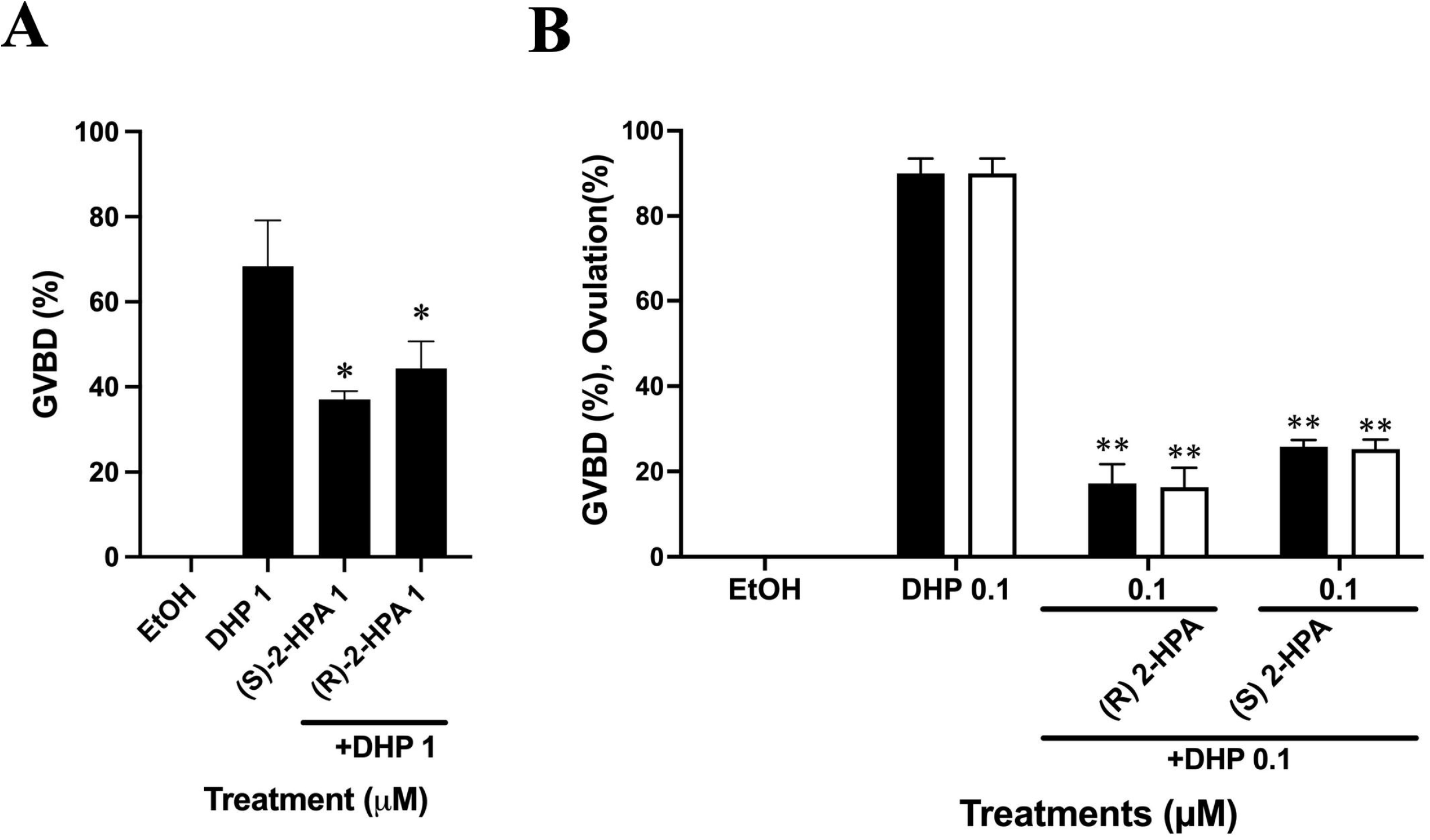
*In vitro* and *in vivo* assay of (S) and (R) types of 2-HPA. (A)The *in vitro* assay results for the (S) and (R) types of 2-HPA are shown. Zebrafish oocytes were used for this low number assay. Each compound was added to the zebrafish Ringer’s solution at a final concentration of 1 μM with 1 μM DHP. After two hours of incubation, the percentage of germinal vesicle breakdown (%GVBD: indicated by the closed column) was determined by scoring the oocytes that became transparent. The assay was performed in triplicate on three fish. Mean of triplicate data is presented with standard deviation. Asterisk indicate significant differences between DHP alone and DHP with 2-HPA treatments (* P≤m 0.05). (B)The *in vivo* assay results for the (S) and (R) types of 2-HPA are shown. Each compound was added to the water at a final concentration of 0.1 μM with 0.1 μM DHP. After four hours of incubation, %GVBD (closed column) and %ovulation (open column) were determined by scoring the oocytes as they became transparent and formed an egg membrane. Three fish per treatment were used. Means of data are presented with standard deviation. Asterisk indicate significant differences between DHP alone and DHP with 2-HPA treatments (** P≤m 0.001).

## Discussion

In this study, a novel natural compound from *Padina arborescence* was identified that interacts with mPRα. In GQD-mPRα binding assay, the compound in peak 2 showed higher affinity than progesterone. Synthetic compounds identified as a major component of Peak2, 2-HPA, showed more higher affinity than Peak2 (Figure 3). Corresponding to this high affinity, 2-HPA showed inhibitory activity against fish oocyte maturation and ovulation at the same concentration of progestin (Figure 5 and 6).

We investigated the structure-activity relationship of 2-HPA analogues to identify the key residues that interact with mPRα. Interestingly, none of the analogues showed activity. The results showed that the hydroxide at position 2 is essential for interaction with mPRα. The length of the carbon chain is also important for binding to mPRα. Thus, it is suggested that the relatively simple structure of 2-HPA is a strict fit for the steroid binding site of mPRα. Conversely, the structural isomers of 2-HPA, (S)-2-HPA and (R)-2-HPA, showed no difference in activity (Figure 6). The results suggest that hydrogens binding to carbon 5 do not contribute to any binding with mPRα.

Treatment with 2-HPA resulted in the prevention of oocyte maturation and ovulation in fish. It can be concluded that the prevention of fish oocyte maturation is due to the binding of 2-HPA to mPRα. Although we cannot exclude the possibility that the inhibition of ovulation is due to the binding of 2-HPA to the nuclear progesterone receptor, which induces ovulation^25^. We believe that this inhibition is due to inhibition of the induction of oocyte maturation, which is the first response in the sequential induction of oocyte maturation and ovulation.

In this study, we have succeeded in identifying the first antagonist of mPRα, the first molecule discovered to mediate a steroid nongenomic action. mPR has five subtypes, and in addition to its function in the reproductive system, which is the subject of this study, mPRα has been implicated in mediating steroid nongenomic actions in the brain and other parts of the body, and in regulating various biological processes. Antagonists of mPRα may be useful for in validating these functions and may also be candidates for new drugs. The identification of novel substances from seaweeds, particularly water-soluble substances, is currently underway. It is highly likely that new antagonists and agonists of mPR will be discovered among these substances. Further elucidation of novel compounds should be encouraged in the future.

## Data availability

The datasets generated and/or analyzed during the current study are available from the corresponding author on reasonable request.

## Methods

### Materials

Chemicals were purchased from companies accordingly: (S)-2-HPA (Combi-Blocks, San Diego, CA); (R)-2-HPA (AMATEK CHEMISTRY, Hong Kong); 2-hydroxybutyric acid (Tokyo Kasei, Japan), valeric acid, 2-hydroxyhexanoic acid, progesterone and 17β-estradiol (Sigma-Aldrich); 1,2-pentanediol, and 3-hydroxyvaleric acid (AdooQ BioScience, Irvine, CA).

### Experimental Fishes

Goldfish were reared and maintained under standard laboratory conditions. The fish used in the experiments were kept in a flow-through aquarium at 20-25°C under a 14-hour light/10-hour dark cycle.

Zebrafish were reared and maintained under standard laboratory conditions. The fish used in the experiments were maintained in a flow-through culture system at 28.5°C under a 14-hour light/10-hour dark cycle.

All fish experiments were approved by the Institutional Ethics Committee of Shizuoka University, Japan (approval numbers 2022F-3 and 2023F-9), and the guidelines for the use of animals established by this committee were strictly adhered to.

### Separation of components in *Padina* secretions by HPLC

Sampling and HPLC purification were carried out as previously described ^24^. Two peaks (Peak1 and 2) were obtained in the final purification step. The Peak2 fraction was dried using a freeze dryer (FDU-810, EYELA). The dried compounds were dissolved in ethanol, which was used for the GQD-hmPRα binding assay and for assays of physiological activities.

### Identification of the major component in the Peak2 from *Padina* secretions by NMR experiment

The Peak2 obtained in the HPLC separation was analyzed by NMR. The fraction of Peak2 was lyophilized and then dissolved in 500 μL of CDCl_3_. The NMR spectra including ^1^H, ^13^C, DQF-COSY, TOCSY, HMQC and HMBC were measured using a JNM-ECZ500R spectrometer (JEOL, Tokyo, Japan), according to the manufacturer’s instructions.

### Evaluation of the specific binding of 2-HPA and its analogues to mPRα

The binding properties of 2-HPA and its analogues to mPRα were evaluated by GQD-hmPRα according to the method described previously^21^.

### *In vitro* goldfish oocyte maturation assay

An *in vitro* oocyte maturation assay was performed in goldfish as described previously^16^.

### *In vitro and in vivo* oocyte maturation assay in zebrafish

An *in vitro and in vivo* zebrafish oocyte maturation and ovulation assay was performed as described previously^15,18^.

## Statistical Analysis

Results were represented as Mean ± SD (Standard Deviation). Data significance values were calculated by using paired t-test. All analytical data and graphs were prepared on GraphPad Prism software version 9.4.0 for Mac OS (GraphPad Software, San Diego, California, USA)

## Acknowledgments

We thank Mr. Daijiro Tokumoto for cultivation of goldfish. This study was supported by Grants-in-Aid for Scientific Research in Priority Areas from the Ministry of Education, Culture, Sports, Science and JSPS KAKENHI Grant Number 20K06719 and 23K05830 (to TT).

## Author contributions statement

MTA and MA performed the purification and the in vitro and in vivo assay. MTA analyzed the data and drafted the manuscript. MSJ conducted GQD-mPR binding assay. MR, MH and MFH collected and cultured *Padina*. SA performed in vitro and in vivo assay. SK performed NMR analysis. TT participated in the study design, supervised the study and wrote the paper. All authors read and approved the final manuscript.

## Additional information

Competing interests: The authors declare no competing interests.

