## Supplemental Figures for "Discovery of specific activity of 2-HPA acting on the membrane progestin receptor α (paqr7) by purification of natural products from the marine algae *Padina*"

Table. S1. Chemical shift values of 2-HPA in the Peak2

| Position | δH | δC |
|---|---|---|
| 1 | | 169.0 |
| 2 | 4.06 | 64.7 |
| 3 | 1.59 | 30.5 |
| 4 | 1.35 | 19.0 |
| 5 | 0.91 | 13.6 |

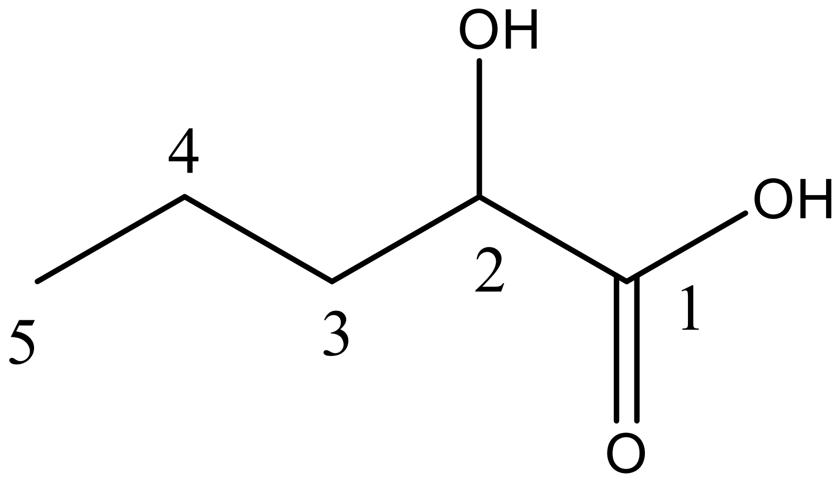

Fig. S1. $^{1}H$ NMR spectrum of the Peak2 in $CDCl_3$

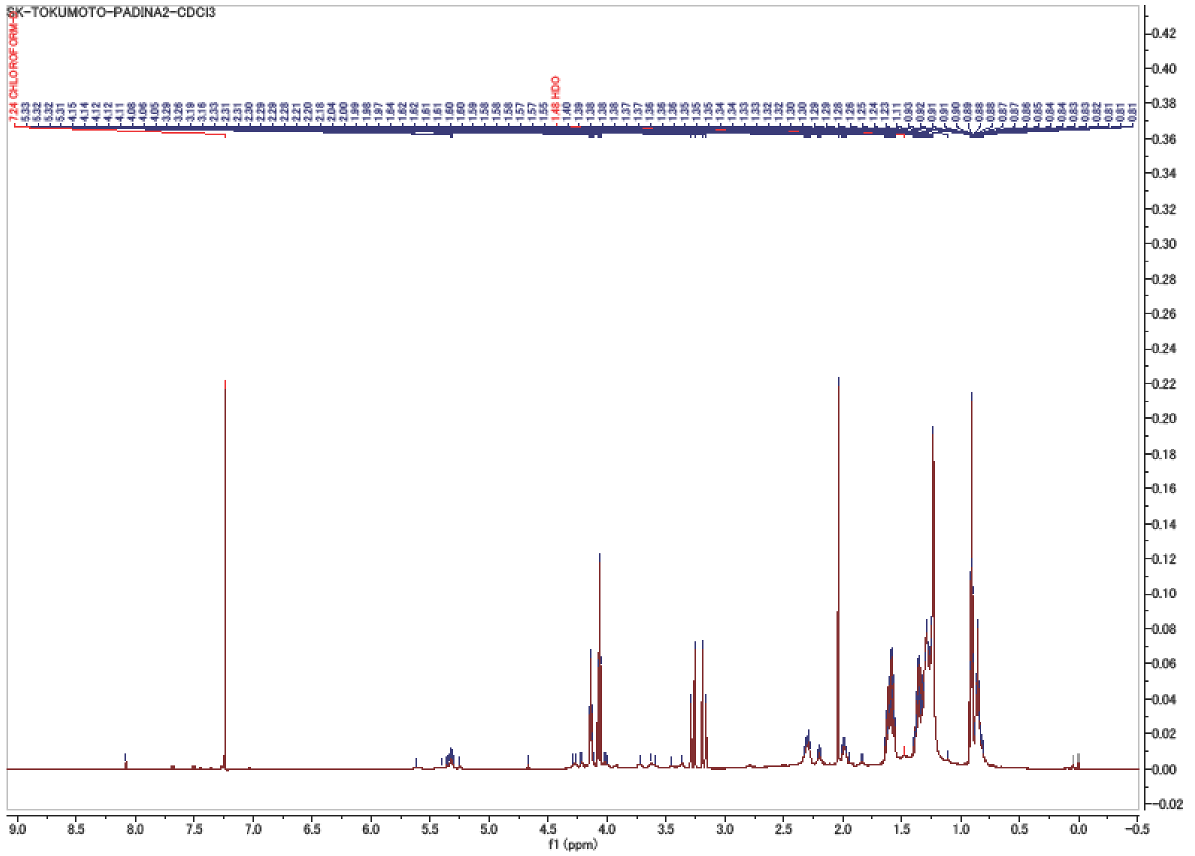

Fig. S2. $^{13}C$ NMR spectrum of the Peak2 in $CDCl_3$

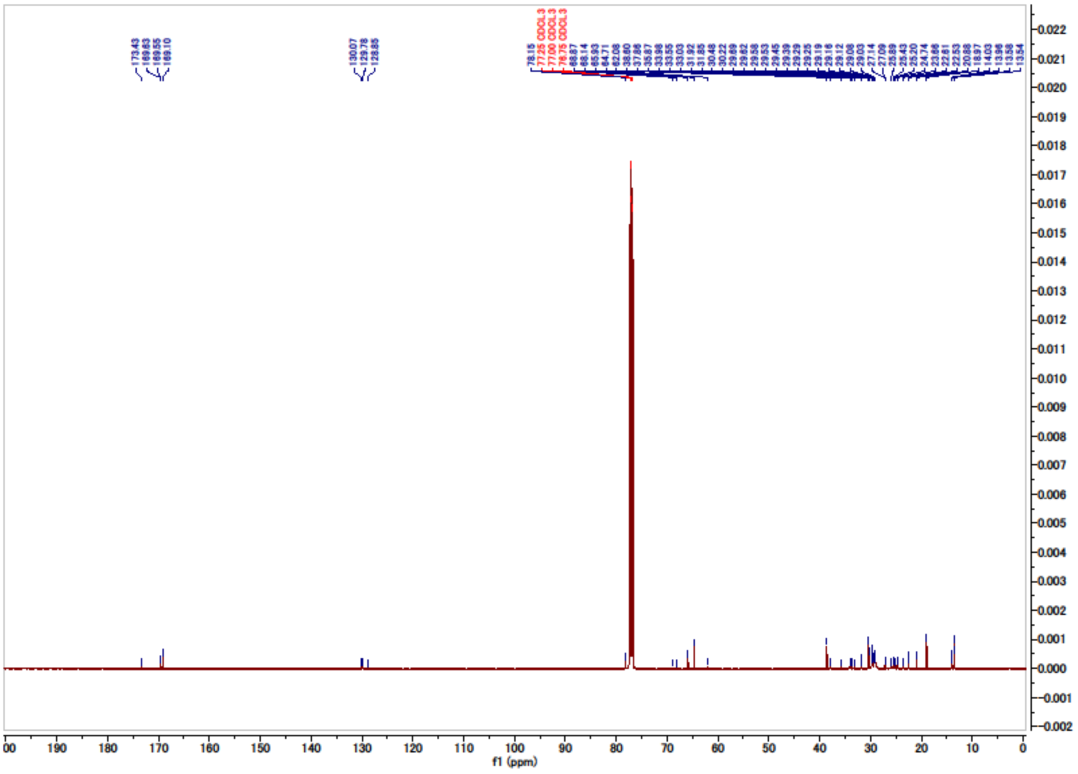

Fig. S3. DQF-COSY spectrum of the Peak2 in $CDCl_3$

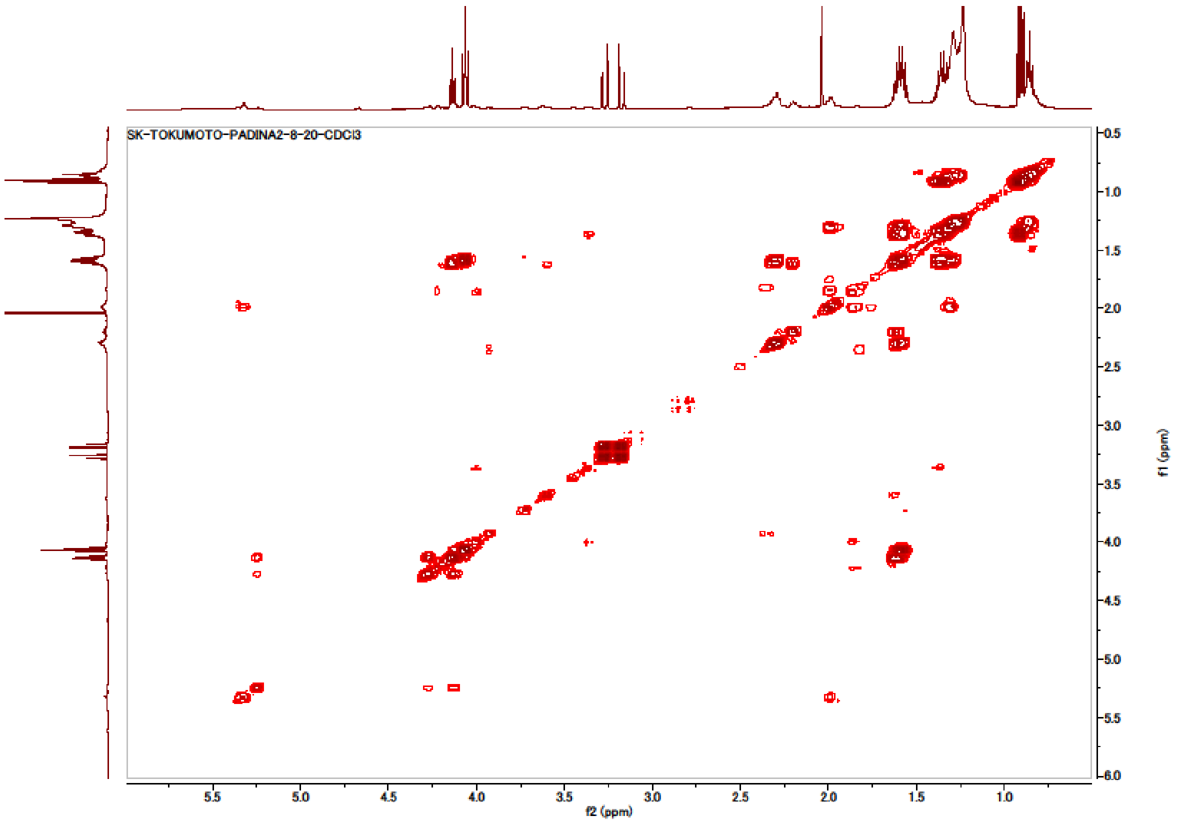

Fig. S4. TOCSY spectrum of the Peak2 in $CDCl_3$

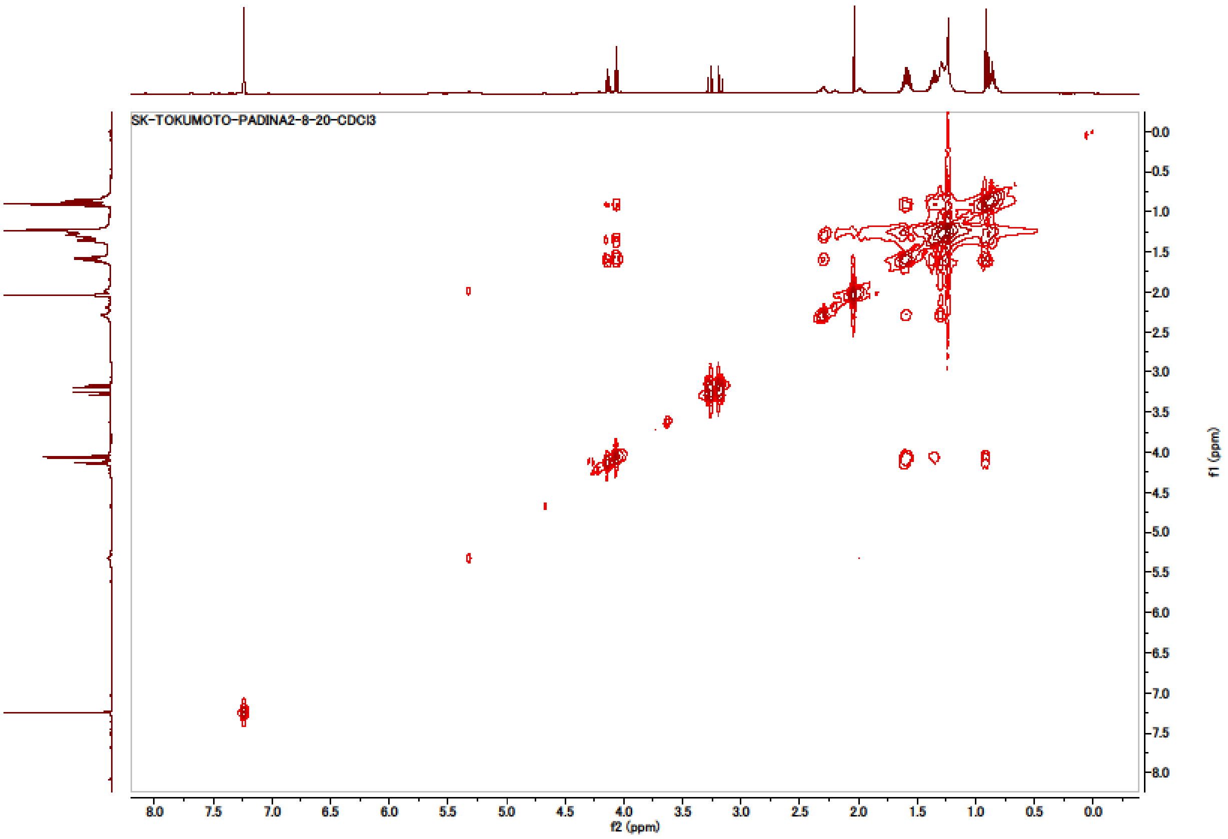

Fig. S5. HMQC spectrum of the Peak2 k in $CDCl_3$

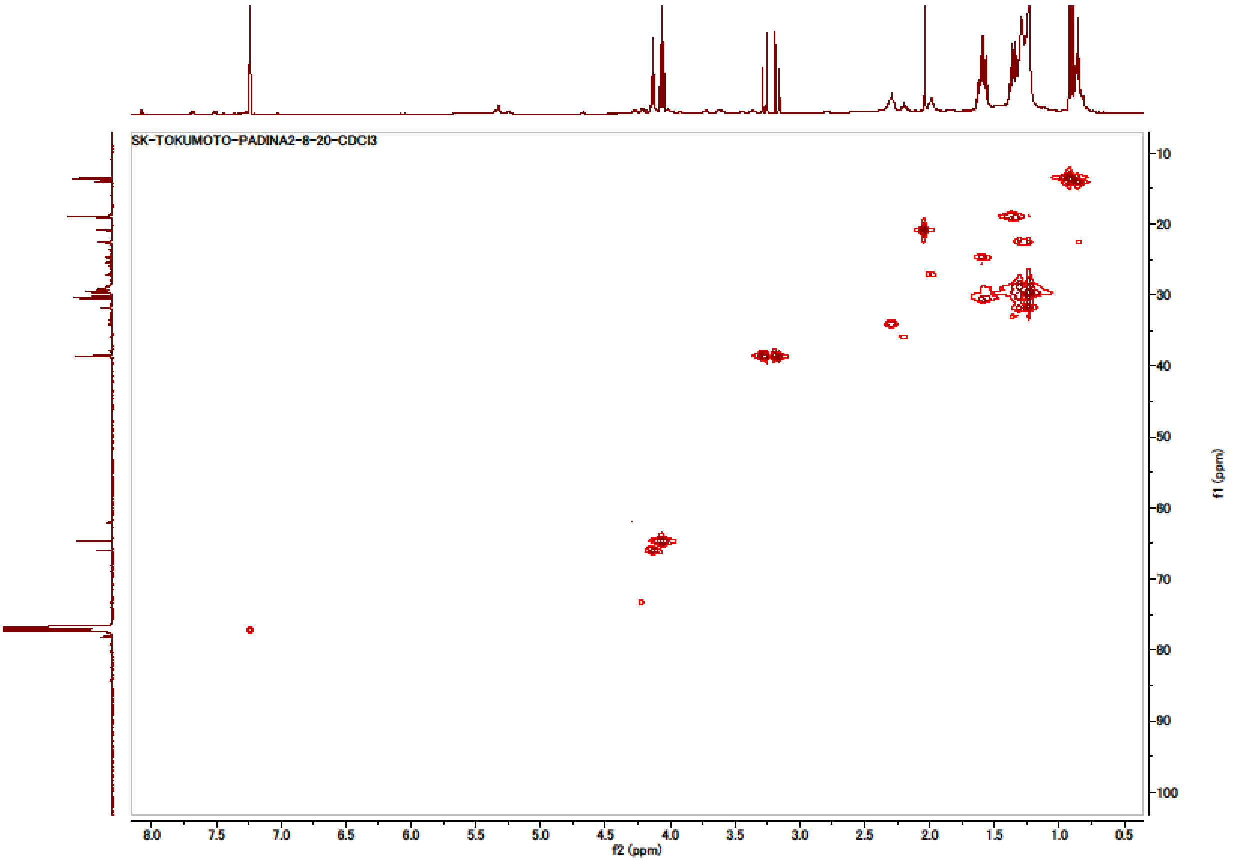

Fig. S6. HMBC spectrum of the Peak2 in $CDCl_3$

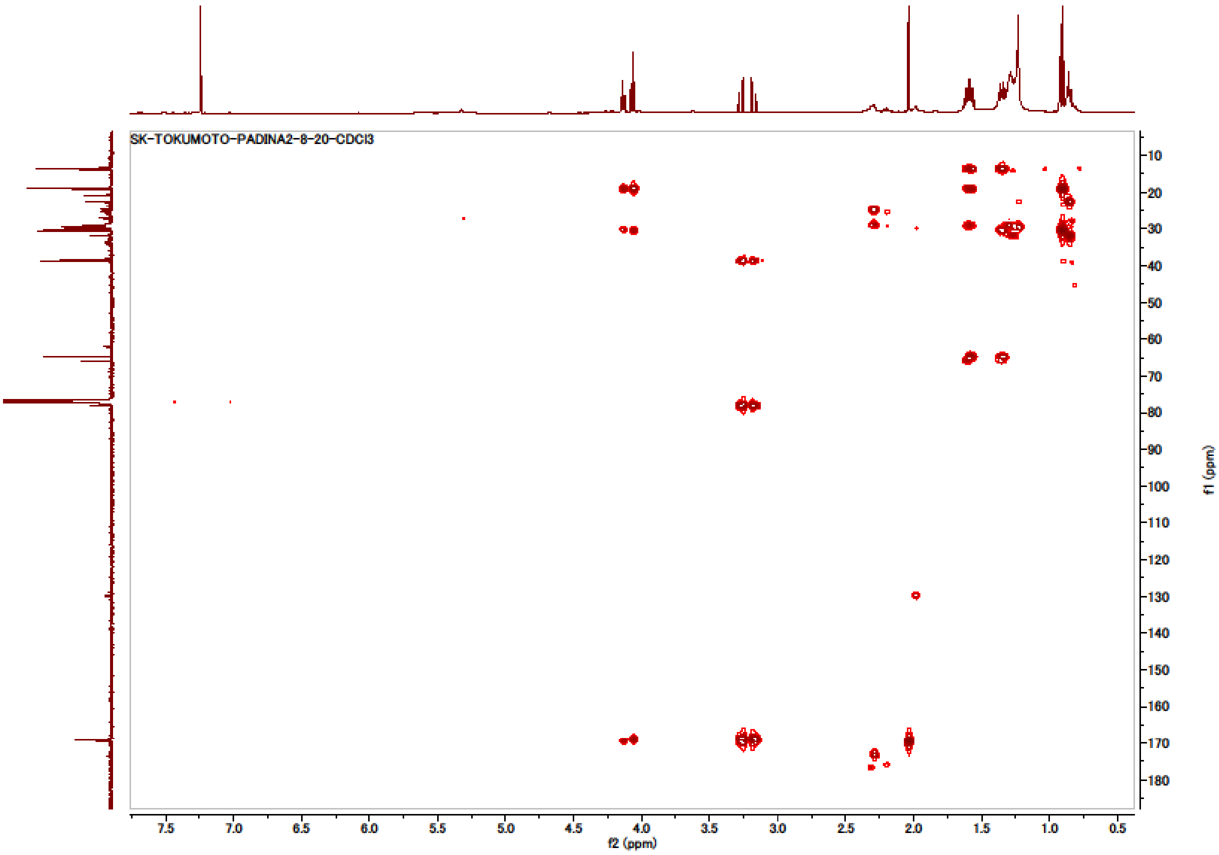
